# Variation in pigmentation gene expression is associated with distinct aposematic color morphs in the poison frog, *Dendrobates auratus*

**DOI:** 10.1101/445684

**Authors:** Adam M. M. Stuckert, Emily Moore, Kaitlin P. Coyle, Ian Davison, Matthew D. MacManes, Reade Roberts, Kyle Summers

## Abstract

Color and pattern phenotypes have clear implications for survival and reproduction in many species. However, the mechanisms that produce this coloration are still poorly characterized, especially at the genomic level. Here we have taken a transcriptomics-based approach to elucidate the underlying genetic mechanisms affecting color and pattern in a highly polytypic poison frog. We sequenced RNA from the skin from four different color morphs during the final stage of metamorphosis and assembled a *de novo* transcriptome. We then investigated differential gene expression, with an emphasis on examining candidate color genes from other taxa. Overall, we found differential expression of a suite of genes that control melanogenesis, melanocyte differentiation, and melanocyte proliferation (e.g., *tyrpl*, *lefl*, *leol*, and *mitf*) as well as several differentially expressed genes involved in purine synthesis and iridophore development (e.g., *arfgap1*, *arfgap2*, *airc*, and *gairt*). Our results provide evidence that several gene networks known to affect color and pattern in vertebrates play a role in color and pattern variation in this species of poison frog.

## Introduction

Color and pattern phenotypes have long been of interest to both naturalists and evolutionary biologists (Bates 1862; Müller 1879). Part of this interest derives from the association of this phenome with selective pressures like mate choice (Kokko et al. 2002) and predation (Ruxton et al. 2004). Species with morphological phenotypes directly tied to survival and reproduction provide excellent opportunities to study the genetic underpinnings of color and pattern, precisely because these phenotypes are so obviously linked to survival.

Aposematic species rely on color and pattern to warn predators, but in many cases these color and pattern phenotypes are extremely variable, often changing over short geographic distances or even exhibiting polymorphism within populations (Brown et al. 2011; Merrill et al. 2015). Theory has long predicted that aposematic species should be monomorphic because predators learn a common signal, and thus aposematic individuals with a different phenotype should be selected against (Müller 1879; Mallet and Joron 1999). While predator variation and drift alone may be sufficient to create phenotypic variation, a variety of alternative selective pressures can act on the aposematic signal to produce and maintain this variety (reviewed in Briolat et al. 2018).

Research on the production of color and pattern early in life in polytypic species (those that vary in discrete phenotypes over geographical space) has been limited, especially in vertebrates. Differences in color and pattern in some highly variable aposematic species seem to be determined by a small number of loci (Martin et al. 2012; Supple et al. 2013; Kunte et al. 2014; Vestergaard et al. 2015). However, the majority of the research on the underlying genetic architecture associated with varied color and patterns in aposematic species has been done in the Neotropical butterflies of the genus *Heliconius*. While this work has been highly informative, it remains unclear whether these trends are generally applicable to other systems, including in vertebrates.

Many of the Neotropical poison frogs (family Dendrobatidae) exhibit substantial polytypism throughout their range (Summers et al. 2003; Brown et al. 2011). Despite being one of the better characterized groups of aposematic species, our knowledge of the mechanisms of color production in this family is quite limited. In addition, there is little information on the genetics of color pattern in amphibians generally. While modern genomic approaches, especially high-throughput sequencing, have recently provided extensive insights into the genes underlying color pattern variation in fish (Diepeveen and Salzburger 2011; Ahi and Sefc 2017), reptiles (Saenko et al. 2013), birds (Ekblom et al. 2012) and mammals (Gene et al. 2001; Bennett and Lamoreux 2003; Bauer et al. 2009), there have been few genomic studies of the genetic basis of color patterns in amphibians. This is in part because amphibian genomes are often large and repetitive. For example the strawberry poison frog (*Oophaga pumilio*) has a large genome (6.7 Gb) which is over two-thirds repeat elements (Rogers et al. 2018). The dearth of amphibian data is an important gap in our knowledge of the genomics of color and pattern evolution, and the genetic and biochemical pathways underlying color pattern variation across vertebrates.

Amphibians exhibit quite varied colors and patterns, and these are linked to the three structural chromatophore types (melanophores, iridophores, and xanthophores) and the pigments and structural elements found within them (e.g. melanins, pteridines and guanine platelets; Mills & Patterson 2009). Melanophores and the melanin pigments they contain are responsible for producing dark coloration, particularly browns and blacks, and are also critical to the production of darker green coloration (Duellman and Trueb 1986). Blue and green coloration in amphibians is generally produced by reflectance from structural elements in iridophores (Bagnara et al. 2007). Iridophores contain guanine crystals arranged into platelets that reflect particular wavelengths of light, depending on platelet size, shape, orientation and distribution (Ziegler 2003; Bagnara et al. 2007; Saenko et al. 2013). Generally speaking, thicker and more dispersed platelets reflect longer wavelengths of light (Saenko et al. 2013). Combinations of iridophores and xanthophores or erythropores containing carotenoids or pteridines (respectively) can produce a wide diversity of colors (Saenko et al. 2013). Xanthophores are thought to be largely responsible for the production yellows, oranges, and reds in amphibians. The precise coloration exhibited is linked to the presence of various pigments such as pteridines and carotenoids that absorb different wavelengths of light (Duellman and Trueb 1986).

In order to better understand the genetic mechanisms affecting the development of color and pattern, we examined four different captive bred color morphs of the green-and-black poison frog (*Dendrobates auratus*). We used an RNA sequencing approach to examine gene expression and characterize the skin transcriptome of this species. In addition to assembling a *de novo* skin transcriptome of a species from a group with few genomic resources, we compared differential gene expression between color morphs. We focused on differential gene expression in a set of *a priori* candidate genes that are known to affect color and pattern in a variety of different taxa. Finally, we examined gene ontology and gene overrepresentation of our dataset. These data will provide useful genomic and candidate gene resources to the community, as well as a starting point for other genomic studies in both amphibians and other aposematic species.

## Methods

### Color morphs

Captive bred *Dendrobates auratus* were obtained from Understory Enterprises, LLC. Four distinct morphs were used in this study; the San Felix and super blue morphs both have a brown dorsum, with the former having green spotting, and the latter typically having light blue markings (often circular in shape), sporadically distributed across the dorsum. The microspot morph has a greenish-blue dorsum with small brownish-black splotches across the dorsum. Finally, the blue-black morph has a dark black dorsum with blue markings scattered across the dorsum that are typically long and almost linear. Photographs of frogs from these morphs in captivity are found in Figure 1. We note that the breeding stock of these different morphs, while originally derived from different populations in Central America, have been bred in captivity for many generations. As a result, it is possible that color pattern differences between these morphs in captivity may exceed those generally found in the original populations. Nevertheless, the differences between these morphs are well within the range of variation in this highly variable, polytypic species which ranges from Eastern Panama to Nicaragua.

**Figure 1.**
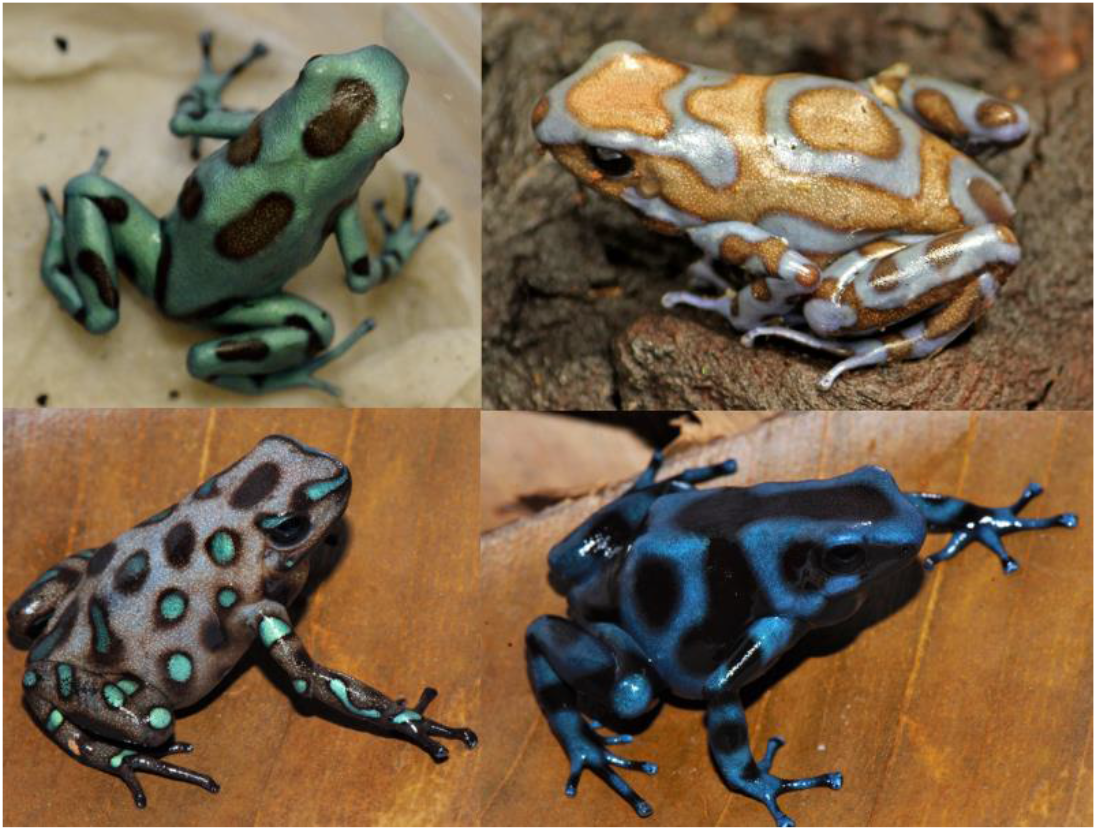
Normative depictions of the four captive morphs used in this study. Color morphs clockwise from top left: microspot, super blue, blue and black, San Felix. Microspot and super blue photographs courtesy of ID, blue-black and San Felix photos were provided by Mark Pepper at Understory Enterprises, LLC.

### Sample collection

Frogs were maintained in pairs in 10 gallon tanks with coconut shell hides and petri dishes were placed under the coconut hides to provide a location for females to oviposit. Egg clutches were pulled just prior to hatching and tadpoles were raised individually in ~100 mL of water. Tadpoles were fed fish flakes three times a week, and their water was changed twice a week. Froglets were sacrificed during the final stages of aquatic life (Gosner stages 41-43; Gosner 1960). At this point, froglets had both hind limbs and at least one forelimb exposed. These froglets had color and pattern elements at this time, but pattern differentiation and color production is still actively occurring during metamorphosis and afterwards. After euthanasia, whole specimens (n = 3 per morph) were placed in RNAlater (Qiagen) for 24 hours, prior to storage in liquid nitrogen. We then did a dorsal bisection of each frog’s skin, and prepared half of the skin for RNA extraction.

RNA was extracted from each bisected dorsal skin sample using a hybrid Trizol (Ambion) and RNeasy spin column (Qiagen) method and total RNA quality was assayed using the Bioanalyzer 2100 (Agilent). Messenger RNA (mRNA) was isolated from total RNA with Dynabeads Oligo(dT)_25_ (Ambion) for use in the preparation of uniquely-barcoded, strand-specific directional sequencing libraries with a 500bp insert size (NEBNext Ultra Directional RNA Library Prep Kit for Illumina, New England Biosystems). Libraries were placed into a single multiplexed pool for 300 bp, paired end sequencing on the Illumina MiSeq. Each sample had a total of 2-5.8 million reads.

### Transcriptome assembly

We randomly chose one individual per morph type and assembled this individual’s transcriptome. First, we aggressively removed adaptors and did a gentle quality trimming using trimmomatic version 0.36 (Bolger et al. 2014). We then implemented read error correction using RCorrector version 1.01 (Song and Florea 2015) and assembled the transcriptome using the Oyster River Protocol version 1.1.1 (MacManes 2018). The Oyster River Protocol (MacManes 2018) assembles a transcriptome with a series of different transcriptome assemblers and kmer lengths, ultimately merging them into a single transcriptome. Transcriptomes were assembled using Trinity version 2.4.0 (Haas et al. 2014), two independent runs of SPAdes assembler version 3.11 with kmer lengths of 55 and 75 (Bankevich et al. 2012), and lastly Shannon version 0.0.2 with a kmer length of 75 (Kannan et al. 2016). The four transcriptomes were then merged together using OrthoFuser (MacManes 2018). Transcriptome quality was assessed using BUSCO version 3.0.1 against the eukaryote database (Simão et al. 2015) and TransRate 1.0.3 (Smith-Unna et al. 2016). BUSCO evaluates the genic content of the assembly by comparing the transcriptome to a database of highly conserved genes. Transrate contig scores evaluate the structural integrity of the assembly, and provide measures of accurate, completeness, and redundancy. We then compared the assembled, merged transcriptome to the full dataset (every read in our dataset concatenated together) by using BUSCO and TransRate.

### Downstream analyses

We annotated our transcriptome using the peptide databases corresponding to frog genomes for *Xenopus tropicalis (NCBI Resource Coordinators 2016), Nanorana parkeri (Sun et al. 2015)*, and *Rana catesbeiana (Hammond et al. 2017)* as well as the UniRef90 database (Bateman et al. 2017) using Diamond version 0.9.10 (Buchfink et al. 2015) and an e-value cutoff of 0.001. We then pseudo-aligned reads from each sample using Kallisto version 0.43.0 (Bray et al. 2016) and examined differential expression of transcripts in R version 3.4.2 (R Core Team 2017) using Sleuth version 0.29.0 (Pimentel et al. 2017). Differential expression was analyzed by performing a likelihood ratio test comparing a model with color morph as a factor to a simplified, null model of the overall data, essentially testing for differences in expression patterns between any of the four morphs. In addition to examining overall differential expression between morphs, we examined differential expression in an *a priori* group of candidate color genes (see supplemental table 1). We used PANTHER (Mi et al. 2017) to quantify the distribution of differentially expressed genes annotated to *Xenopus tropicalis* into biological processes, molecular functions, and cellular components.

### Data and analyses availability

All read data are archived with the European Nucleotide Archive project PRJEB25664 (embargoed until paper acceptance). Code for transcriptome assembly, annotation, and downstream analyses are all available on GitHub (https://github.com/AdamStuckert/Dendrobates_auratus_transcriptome). Further, our candidate color genes are available in Supplementary Table 1, and for the purposes of review, our assembled transcriptome is publicly available (Stuckert 2018; http://doi.org/10.5281/zenodo.1443579).

## Results

### Transcriptome assembly

After conducting the Oyster River Protocol for one random individual per color morph and merging them together, we were left with a large transcriptome containing 597,697 transcripts. We examined the BUSCO and transrate scores for each morph’s transcriptome, as well as for the transcriptome created by orthomerging these four assemblies (Table 1). BUSCO and transrate scores were computed using the full, cleaned read dataset from all samples. Given the poor transrate score of our final, merged assembly we selected and used the good contigs from transrate (i.e., those that are accurate, complete, and non-redundant), which had a minimal effect on our overall BUSCO score. In total, our assembly from the good contigs represents 160,613 individual transcripts (the “full assembly” in Table 1). Overall, our annotation to the combined *Xenopus*, *Nanorana*, *Rana*, and UniRef90 peptide databases yielded 76,432 annotated transcripts (47.5% of our transcriptome).

**Table 1.** Assembly metrics for each of our assembled transcriptomes. Metrics for the full assembly were calculated using the full, cleaned dataset. BUSCO scores represent the percentage of completion (i.e., 100% is an entirely complete transcriptome).

|  | Transrate score | Transrate optimal score | BUSCO score |
| --- | --- | --- | --- |
| Blue-black | 0.05446 | 0.40487 | 96.3% |
| Microspot | 0.04833 | 0.35907 | 94.0% |
| San Felix | 0.0556 | 0.35718 | 88.1% |
| Super blue | 0.0521 | 0.38094 | 96.0% |
| Full assembly | 0.01701 | 0.13712 | 95.8% |

### Differential expression and pathways

Our results indicate that there are distinct differences in expression between color morphs (Figure 2). Principal component 1 (37.3% of variation explained) and principal component 2 (21.0% of variation explained). When we tested for differential expression we found a total of 2,845 differentially expressed transcripts among color morphs (1.77% of our transcriptome; Supplementary Table 2). From our list of candidate color genes, we found 58 differentially expressed transcripts (q value < 0.05) associated with 41 candidate color genes in total (see Table 2 and Figures 6 and 7). Many of these genes are involved in typical vertebrate pigmentation pathways, which we highlight in Figure 8. In our analyses of gene function using all differentially expressed genes in PANTHER, we found that most of these genes were associated with either metabolic or cellular processes (Figure 3). Similarly, most of these genes contributed to either cell part or organelle cellular components (Figure 4). The molecular function was heavily skewed towards catalytic activity and binding, both of which are likely a result of the huge developmental reorganization involved in metamorphosis (Figure 5).

**Figure 2.**
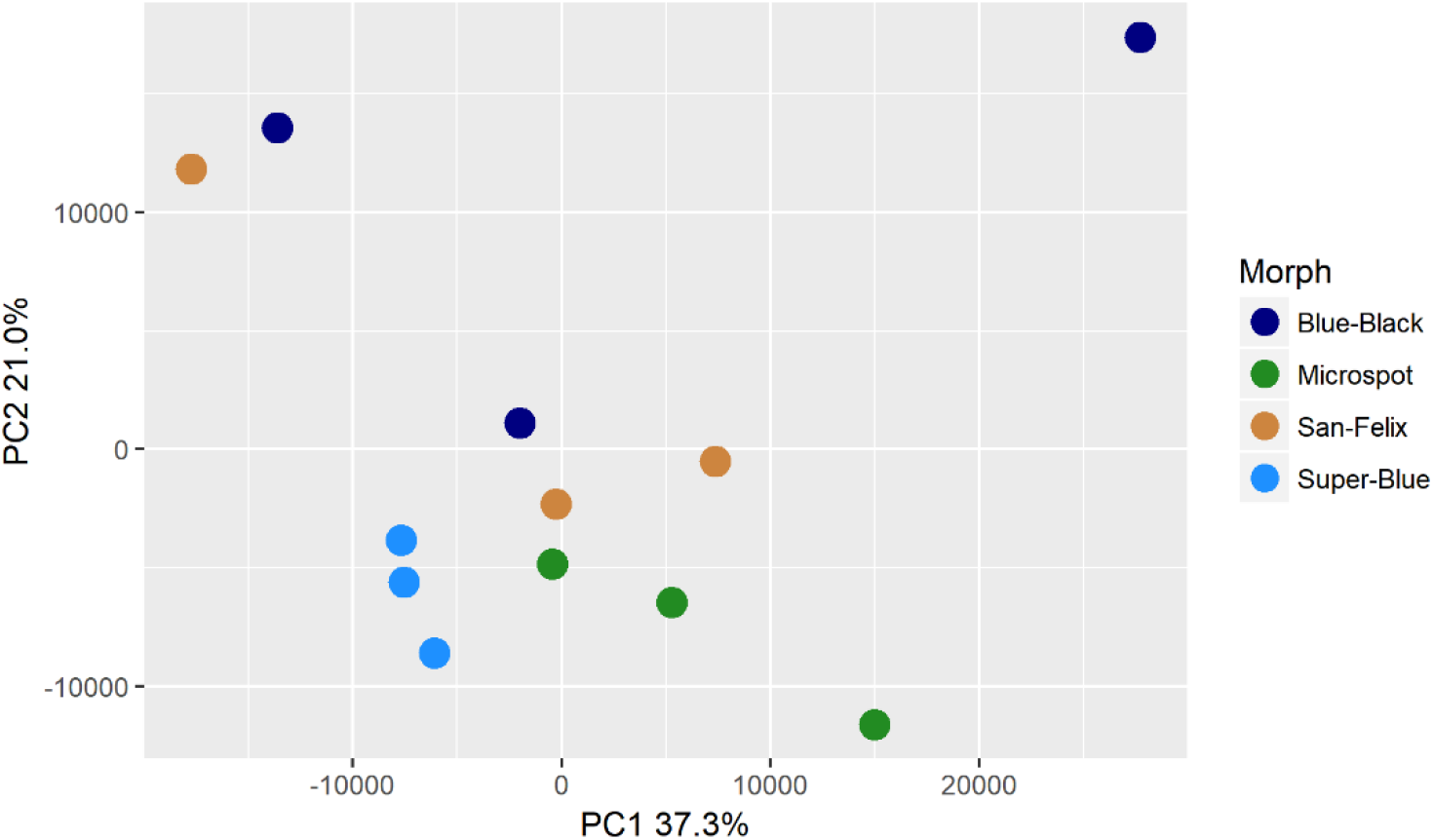
Principal component analysis indicating general within-morph similarity in transcript abundance within our dataset. PCA computation was normalized as transcripts per million. Each dot indicates one individual and the percentage of variation explained by the axes are presented.

**Table 2.** Differentially expressed candidate color genes in our *Xenopus* annotation. Parentheses in the gene symbol column indicate the number of transcripts that mapped to a particular gene. The pathway column indicates what color or pattern production pathway this gene is a part of.

| Gene symbol | q value | Pathway | Citation |
| --- | --- | --- | --- |
| adam17 (2) | 0.0163;<br>0.0469 | Melanocyte development | Bennett and Lamoreux 2003 |
| arfgap1 (2) | 0.00362;<br>0.0267 | Putative guanine synthesis in iridophores | Higdon et al. 2013 |
| arfgap3 (4) | 0.00739;<br>0.0000123;<br>0.00132;<br>0.0282 | Putative guanine synthesis in iridophores | Higdon et al. 2013 |
| airc | 0.0126 | Guanine synthesis | Tolstorukov and Efremov 1984; Sychrova et al. 1999 |
| atic | 0.0447 | Guanine synthesis in iridophores | Higdon et al. 2013 |
| atox1 | 0.00124 | Melanogenesis | Hung et al. 1998; Klomp et al. 1997 |
| atp12a | 0.0296 | Melanogenesis | Nelson et al. 2009 |
| bbs2 | 0.0300 | Melanosome transport | Tayeh et al. 2008 |
| bbs5 | 0.0447 | Melanosome transport | Tayeh et al. 2008 |
| bmpr1b | 0.0118 | Inhibits melanogenesis | Yaar et al. 2006 |
| brca1 | 0.0455 | Alters pigmentation, produces piebald appearances in mice | Ludwig et al. 2001; Tonks et al. 2012 |
| ctr9 | 0.0280 | Melanocyte assembly | Akanuma et al. 2007; Nguyen et al. 2010 |
| dera |  | Guanine synthesis in iridophores | Higdon et al. 2013 |
| dio2 (3) | 0.0338;<br>0.0256;<br>0.000866 | Thyroid hormone pathways, tenuous | McMenamin et al. 2014 |
| dtnbp1 (2) | 0.00120;<br>0.0456 | Melanosome biogenesis | Wei 2006 |
| ednrb (2) | 0.0035;<br>0.0005 | Guanine synthesis in iridophores, melanoblast migration | Higdon et al. 2013; Kelsh et al. 2009 |
| egfr (2) | 0.0197;<br>0.000566 | Melanocyte pigmentation and differentiation | Jost et al. 2000; Hirobe 2011 |
| fbxw4 (2) | 0.00268;<br>0.0183 | Melanophore organization | Kawakami et al. 2000; Ahi and Sefc 2017 |
| gart | 0.0000494 | Purine synthesis, affecting iridophores, xanthophores, and melanophores | Ng et al. 2009 |
| gas1 (2) | 0.0264;<br>0.0191 | Guanine synthesis in iridophores | Higdon et al. 2013 |
| gne (2) | 0.00571;<br>0.0361 | Sialic acid pathway | Nie et al. 2016 |
| hps3 | 0.0202 | Melanosome biogenesis | Suzuki et al. 2001 |
| itgb1 (2) | 0.0191;<br>0.0469 | Guanine synthesis in iridophores | Higdon et al. 2013 |
| lef1 | 0.0190 | Melanocyte differentiation and development, melanogenesis | Song et al. 2017 |
| leo1 | 0.0000381 | Melanocyte assembly | Johnson et al. 1995 |
| mitf | 0.0466 | Melanocyte regulation | Levy et al. 2006; Hou and Pavan 2008 |
| mlph | 0.00568 | Melanosome transport | Cirera et al. 2013 |
| mthfd1 | 0.0430 | Purine synthesis | Field et al. 2011 |
| mreg | 0.0156 | Melanosome transport | Wu et al. 2012 |
| notch1 (3) | 0.00681;<br>0.0139;<br>0.0487 | Melanocyte production | Shouwey and Beerman 2008 |
| prtfdc1 | 0.00000672 | Guanine synthesis | Higdon et al. 2013 |
| qdpr | 0.0372 | Guanine and Pteridine synthesis | Xu et al. 2014; Ponzone et al. 2004 |
| qnr-71 (2) | 0.0316;<br>0.0262 | Melanosomal protein | Turque et al. 1996; Planque et al. 1999 |
| rab3d | 0.0321 | Putative guanine synthesis in iridophores | Higdon et al. 2013 |
| rab7a | 0.0319 | Putative guanine synthesis in iridophores | Higdon et al. 2013 |
| rabggta | 0.000864 | Guanine synthesis | Swank et al. 1993 |
| scarb2 | 0.0329 | Putative guanine synthesis in iridophores | Higdon et al. 2013 |
| shroom2 | 0.0142 | Pigment accumulation | Fairbank et al. 2006; Lee et al. 2009 |
| sox9 | 0.0228 | Melanin production | Passeron et al. 2007 |
| tbx15 | 0.00838 | Pigmentation boundaries | Candille et al. 2004 |
| tyrp1 | 0.0200 | Melanogenesis | Rieder et al. 2001 |
| xdh (2) | 0.0346;<br>0.0384 | Pteridine synthesis | Thorsteinsdottir and Frost 1986 |

**Figure 3.**
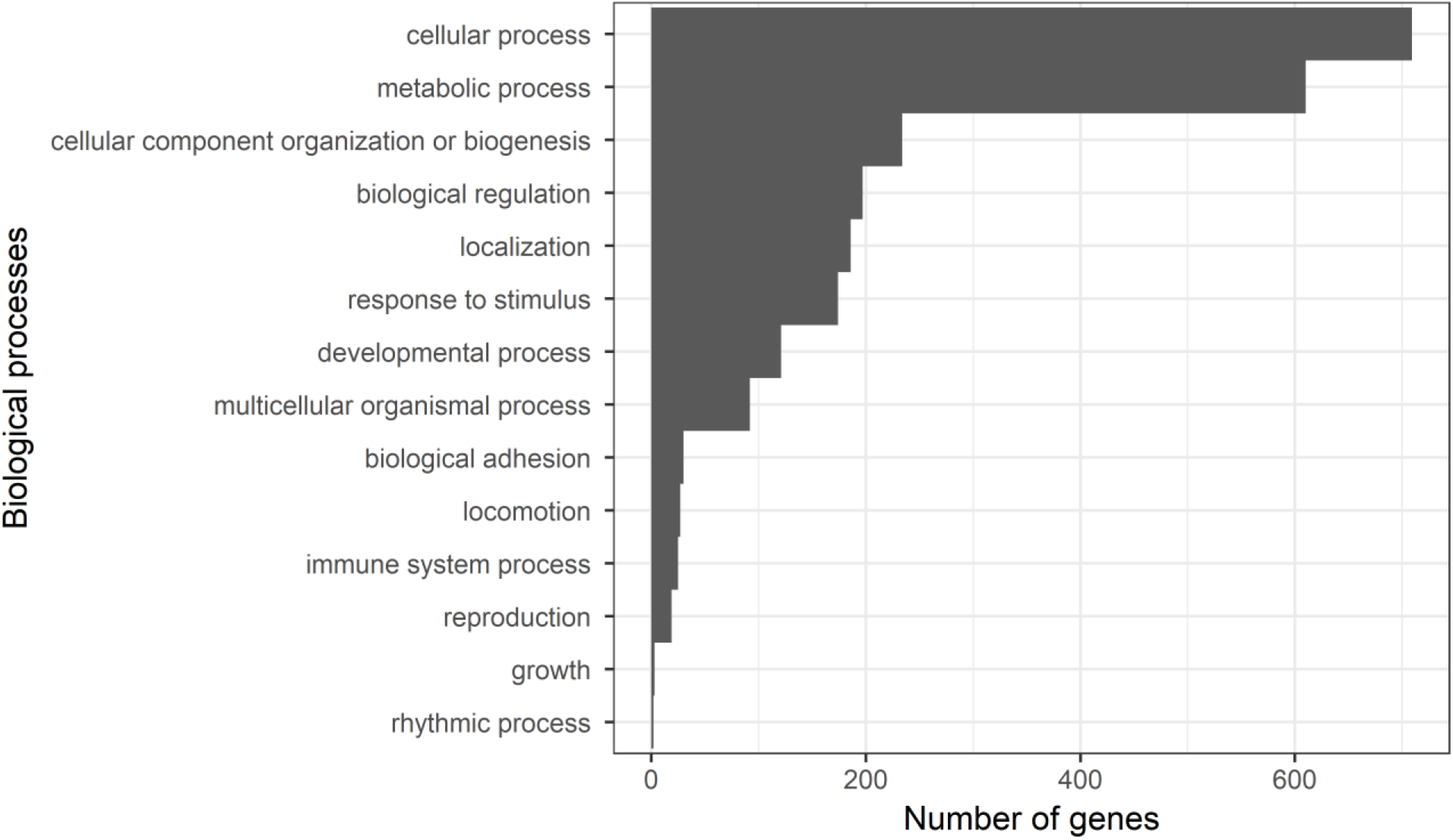
Gene ontology terms from PANTHER. Bars depict the number of genes in each biological process GO category.

**Figure 4.**
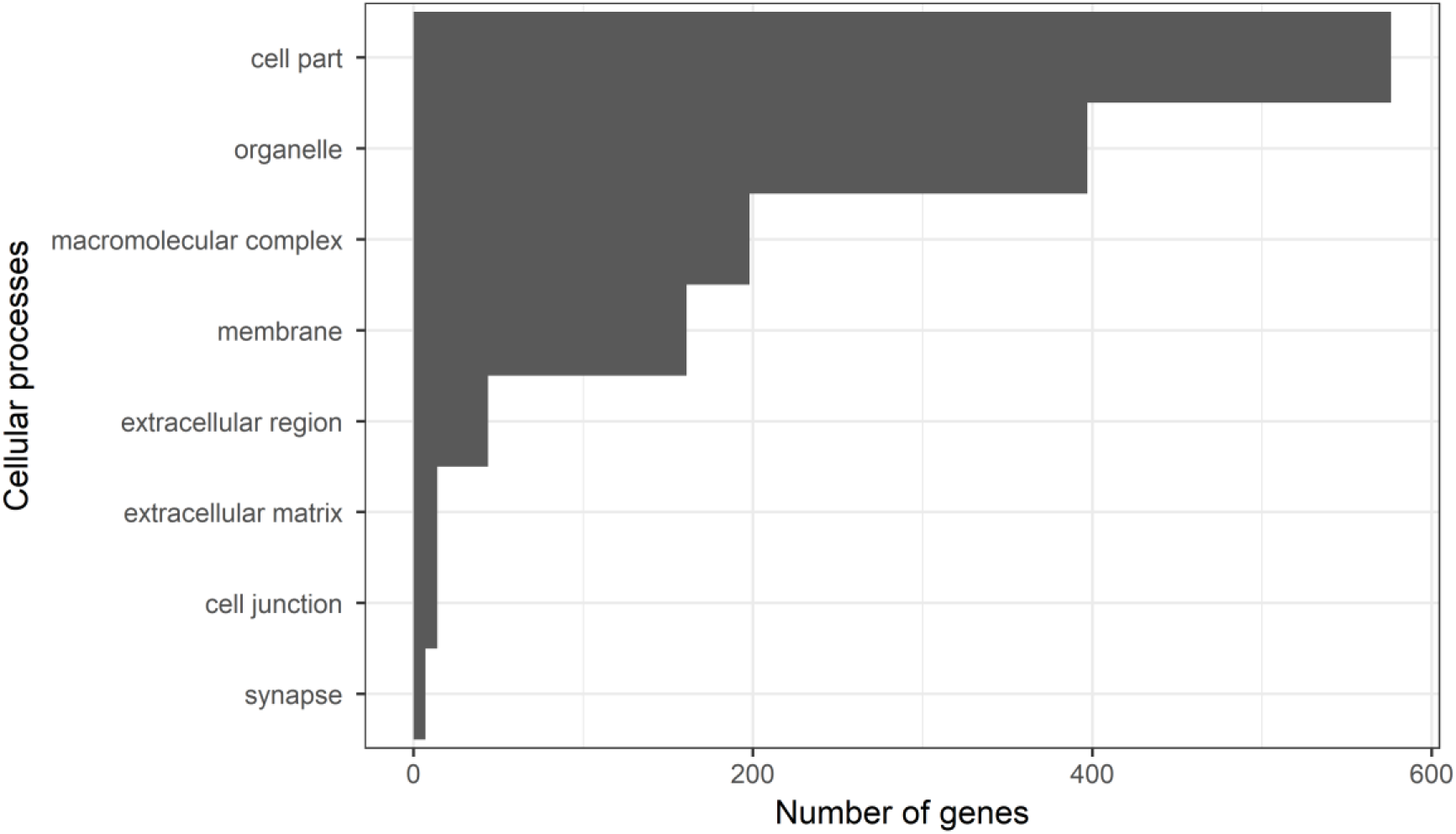
Gene ontology terms from PANTHER. Bars depict the number of genes in each cellular process GO category.

**Figure 5.**
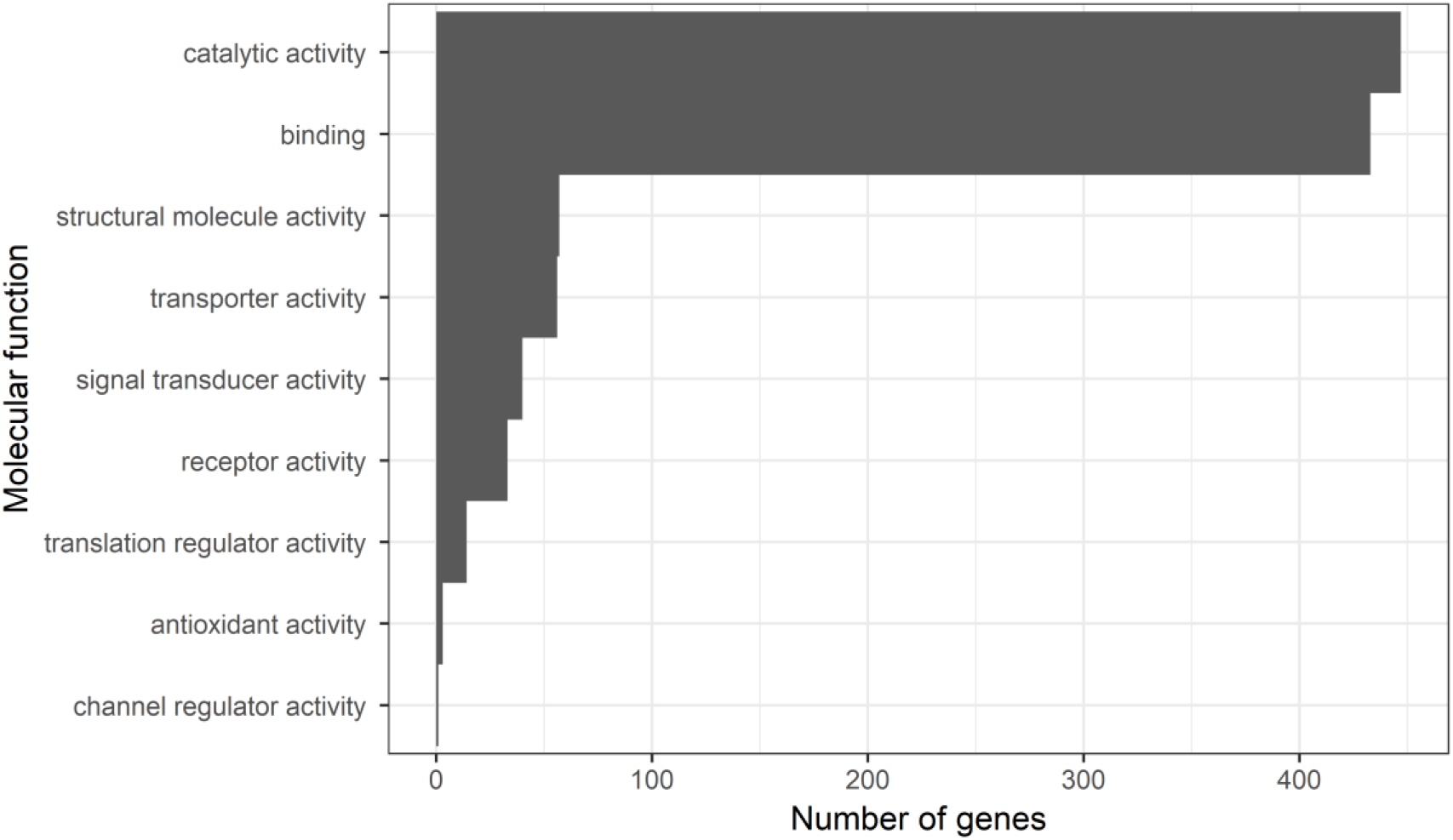
Gene ontology terms from PANTHER. Bars depict the number of genes in each molecular function GO category.

**Figure 6.**
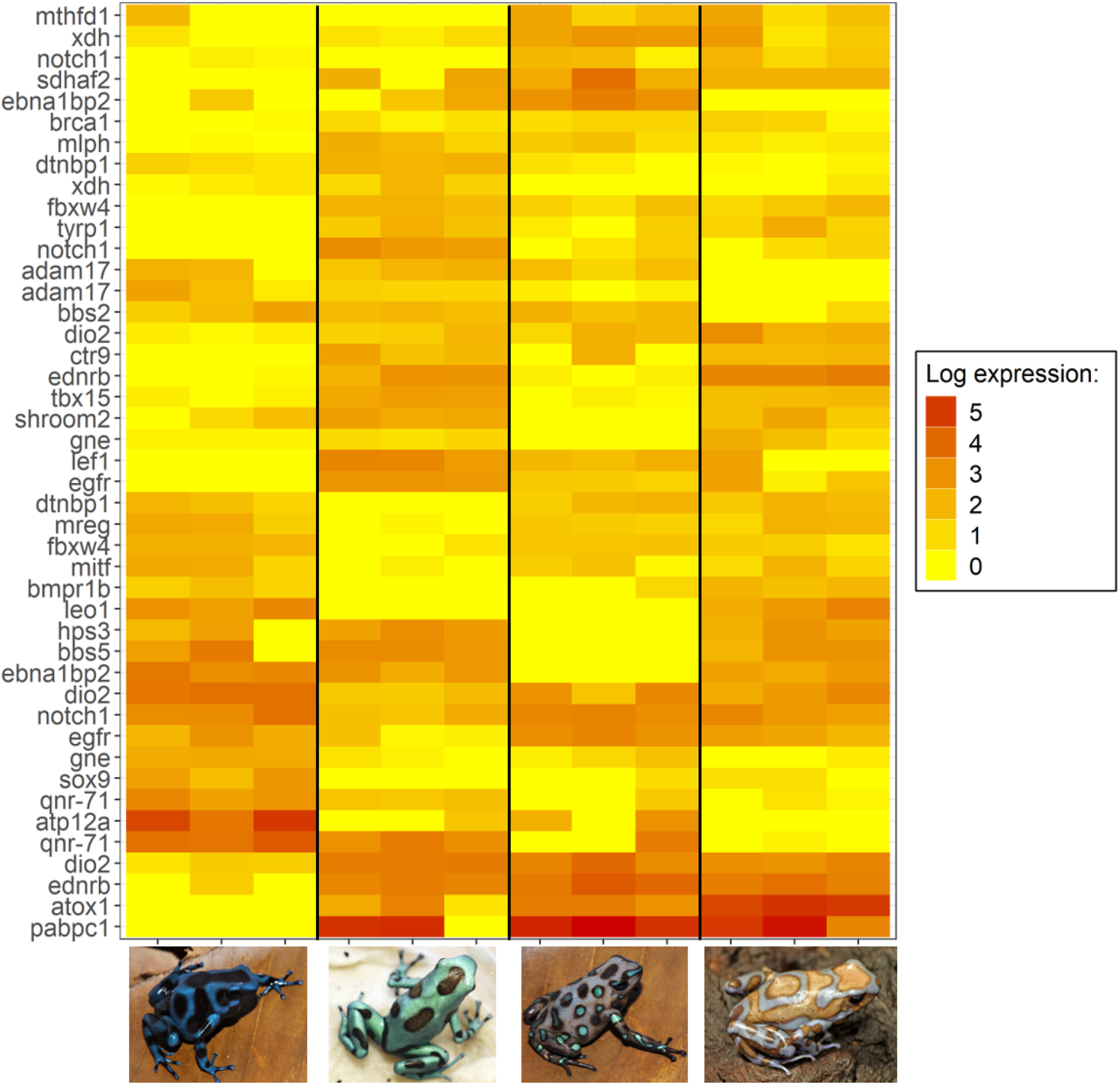
Log-fold expression (transcripts per million) levels of putatively melanin related genes in *Dendrobates auratus*. Each individual is represented on the x-axis, and the y-axis represents expression levels for each transcript that annotated to an melanophore-related gene. Genes represented more than once mapped to multiple transcripts. Expression for this heatmap was calculated using the transcripts per million from Kallisto, to which we added 1 and log transformed the data (i.e., expression = log(transcripts per million + 1).

**Figure 7.**
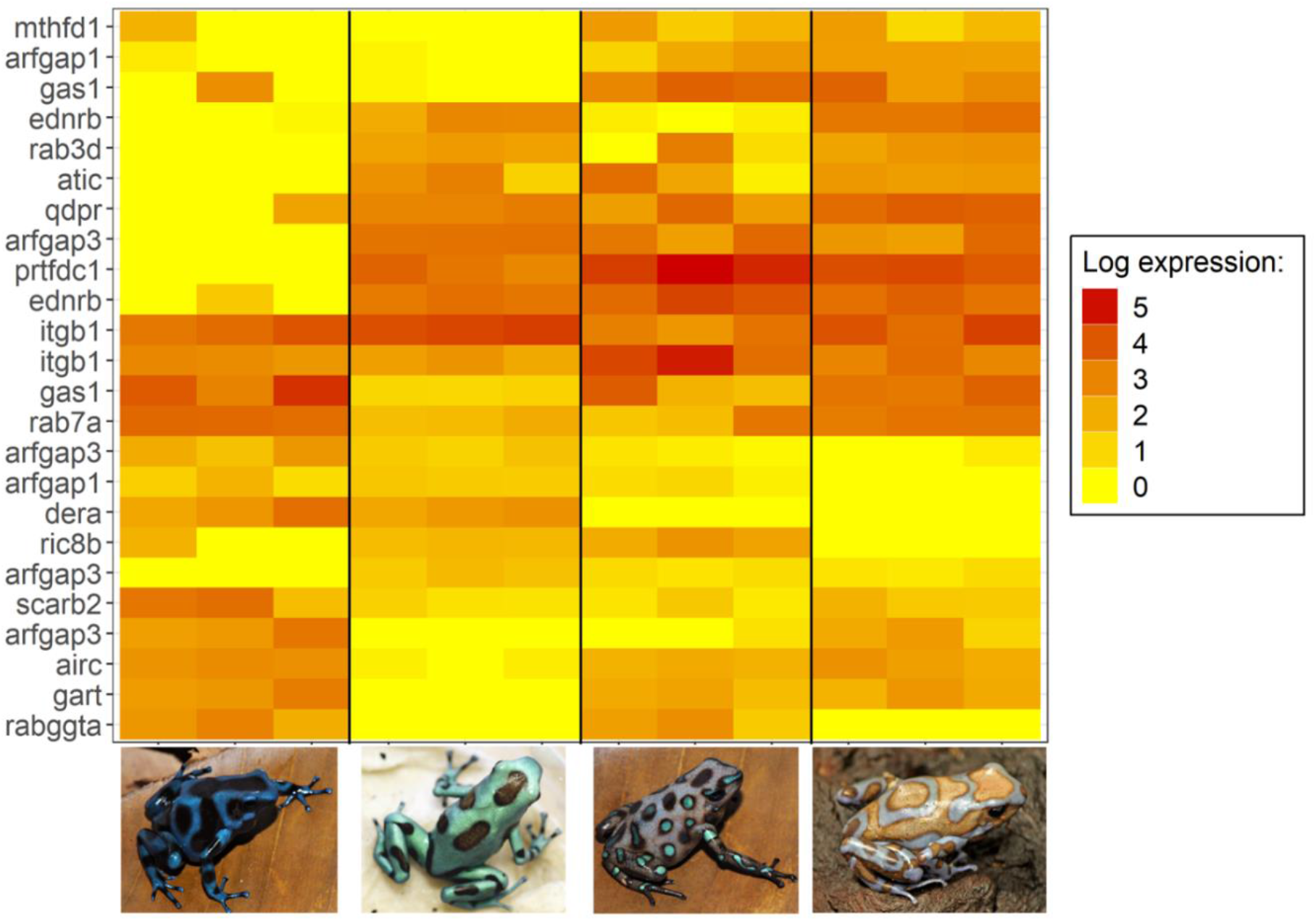
Log-fold expression (transcripts per million) levels of putatively iridophore-related genes in *Dendrobates auratus*. Each individual is represented on the x-axis, and the y-axis represents expression levels for each transcript that annotated to an iridophore-related gene. Genes represented more than once mapped to multiple transcripts. Expression for this heatmap was calculated using the transcripts per million from Kallisto, to which we added 1 and log transformed the data (i.e., expression = log(transcripts per million + 1)).

**Figure 8.**
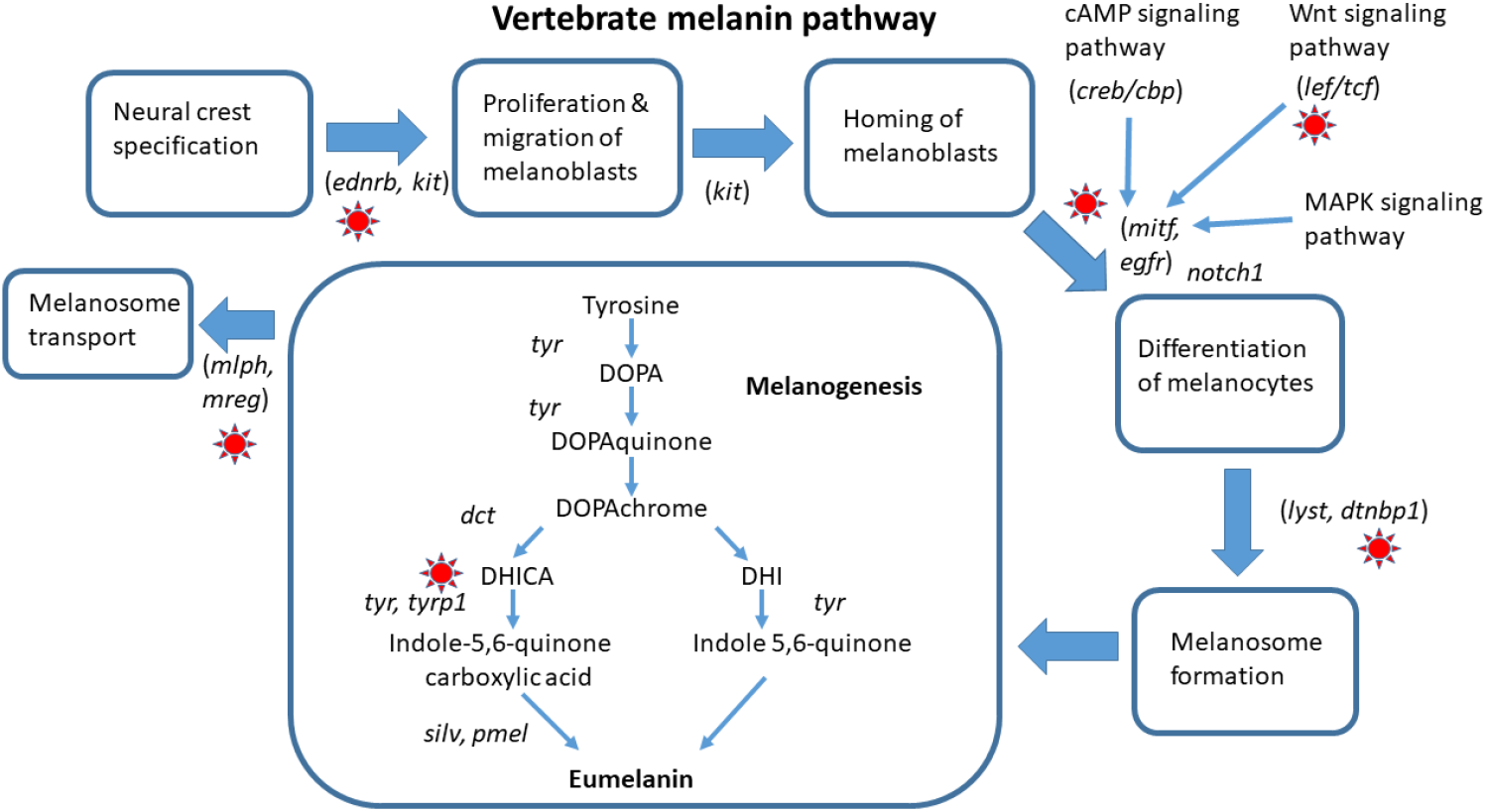
Melanin pigmentation pathway in vertebrates. Here we highlight differentially expressed genes in our dataset with a red sun.

## Discussion

The genetic mechanisms of color variation are poorly known, particularly in amphibians. Here, we address this deficiency by providing some of the first genomic data relevant to color production in amphibians, with a focus on gene expression in the skin during development. Our model system and strategy support the identification of genes likely to regulate color and pattern elements across different morphs of a highly variable species. By combining analyses of differential expression with a targeted search based on an extensive list of candidate genes for developmental control of coloration (approximately 500 genes), we identified multiple genes that were differentially expressed among morphs which have been demonstrated to play important roles in the production of color in other taxa.

We found differential expression of multiple genes in two major suites of color genes, those that influence melanic coloration (black, brown, and grey) and iridophore genes (blue and green coloration). Additionally, we found a few key pteridine pigment genes that are known to influence primarily yellow amphibian coloration that were differentially expressed between morphs. Given that our color morphs had a black versus brown color coupled with either blue or green pattern elements on top of the background, these results seem biologically relevant and indicative of genes that control color and pattern in *Dendrobates auratus*. As a result, we divide our discussion into three main parts, focusing on the genes that influence dark background coloration, purine synthesis, and iridophore biology. We then discuss a few genes that are part of other pathways (e.g. pteridine synthesis), before proposing genes that have yet to be implicated in the production of color but are plausible candidate genes.

### Melanin-related gene expression

Our study frogs have skin with either a black or brown background, both of which are forms of melanic coloration, which provides the basis for contrasting patterns in many vertebrates as well as non-vertebrate taxa (Sköld et al. 2016). Melanin is synthesized from tyrosine in vertebrates, via the action of a set of key enzymes (e.g., tyrosinase, tyrosinase-like protein 1 and 2). We identified a suite of differentially expressed genes that are involved in the production of melanophores and melanin in this study (Figure 6 and 8), many of which have been tied to the production of relatively lighter phenotypes in previous studies. Intriguingly, our results parallel similar findings in *Oophaga histrionica*, a species of poison frog in which mutations in the *mc1r* gene affecting melanogenesis have produced a lighter, more brownish background in some populations (Posso-Terranova and Andrés 2017). In a pattern reminiscent of their results, we found that *mc1r* was only lowly expressed in one super blue frog, and that a variety of other genes linked to lighter phenotypes followed a similar pattern of expression.

For example, many of the differentially expressed color genes in our dataset are active contributors to the tyrosinase pathway (*tyrp1*, *mitf*, *sox9*, *lef1*, *mlph*, *leo1*, *adam17*, *egfr*, *ednrb*). This pathway is enzymatically regulated by tyrosinase as well as other enzymes and cofactors and is key to the production of melanin (Murisier and Beermann 2006). The *tyrp1* enzyme catalyzes several key steps in the melanogenesis pathway in melanosomes (and melanocytes), has been shown to affect coloration in a wide variety of vertebrates (Murisier and Beermann 2006; Braasch et al. 2009), and is important for maintaining the integrity of the melanocytes (Gola et al. 2012). In some mammals *tyrp1* has been shown to change the relative abundances of the pigments pheomelanin and eumelanin, thereby producing an overall lighter phenotype (Videira et al. 2013). Our data mimic this pattern as *tryp1* is not expressed in the blue-black morph, and only expressed at low levels in some San Felix individuals. Pheomelanin has only been identified in the skin of one species of frog (Wolnicka-Glubisz et al. 2012), and it is unclear whether pheomelanin is generally present in ectotherms. Further, mutations in *tyrp1* change melanic phenotypes through different mechanisms in fish (and possibly other ectotherms) than in mammals (Braasch et al. 2009; Cal et al. 2017), and the mechanisms by which *tyrp1* one affects pigmentation in amphibians are still being elucidated.

The *mitf* (microphthalmia-associated transcription factor) locus codes for a transcription factor that plays a dominant role in melanogenesis, and has been called the “master regulator” of melanogenesis (Kawakami and Fisher 2017). In our study, *mitf* expression was lowest in the microspot population, the population with the least melanic coloration, and most highly expressed in the blue-black morph (although it is worth noting that blue and green colors are also influenced by melanin to some degree). The *mitf* locus is, itself, targeted by a suite of transcriptional factors including two which were differentially expressed in our dataset: *sox9* and *lef1*. The *sox9* gene is upregulated during melanocyte differentiation, can promote melanocyte differentiation, and has been demonstrated to be an important melanocytic transcription factor (Cheung and Briscoe 2003). Further, *sox9* is up-regulated in human skin after UVB exposure and has been demonstrated to increase pigmentation. *Sox9* was not expressed in the microspot morph and was only expressed (at a low level) in one San Felix individual. Another important transcription factor is the lymphoid enhancer-binding factor locus (*lef1*), which mediates *Wnt* signaling in the context of melanocyte differentiation and development, with important effects on melanogenesis (Song et al. 2017). Upregulation of this gene has been found to reduce synthesis of the darkest melanic pigment eumelanin, resulting in lighter coloration in mink and other vertebrates (Song et al. 2017). In our study, *lef1* showed very low expression in the blue and black morph, compared to the other three morphs. Comparing the photos of the four morphs (Fig. 1), it can readily be seen that blue and black morph has substantially darker (black) background coloration, compared to the other three, which all have a lighter, brownish background coloration indicating that *lef1* is a likely contributor to the background dorsal coloration between color morphs in *Dendrobates auratus*.

Just as *mitf* is a target of the transcription factors *lef1* and *sox9*, *mitf* targets endothelin receptors, a type of G Protein Coupled Receptor (GPCR). Endothelin receptors mediate several crucial developmental processes, particularly the development of neural crest cell populations (Braasch and Schartl 2014). Three paralogous families of these receptors have been identified in vertebrates: endothelin receptor B1 (*ednrb1*), endothelin receptor B2 (*ednrb2*), and endothelin receptor A (*ednra). Ednrb* is involved in producing the different male color morphs of the Ruff (a sandpiper), and it is only expressed in black males (Ekblom et al. 2012). In our study, *ednrb* is not expressed in the blue-black morph, and only one of the *ednrb* transcripts is expressed in the San Felix morph. Mutations in *ednrb1* and *ednrb2* have been found to affect pigment cell development (especially melanocytes and iridophores) in a variety of vertebrate species (Braasch and Schartl 2014). These receptors show divergent patterns of evolution in the ligand-binding region in African lake cichlids, and appear to have evolved divergently in association with adaptive radiations in this group (Diepeveen and Salzburger 2011). The *ednrb2* (endothelin receptor B2) locus encodes a transmembrane receptor that plays a key role in melanoblast (a precursor cell of the melanocyte) migration (Kelsh et al. 2009). This receptor interacts with the *edn3* ligand. Mutations affecting this ligand/receptor system in *Xenopus* affect pigment cell development (Kawasaki-Nishihara et al. 2011).

The *leol* (LEO1 Homolog) and *ctr9* (CTR9 Homolog) loci are both components of the yeast polymerase-associated factor 1 (*Paf1*) complex, which affects the development of the heart, ears and neural crest cells in zebrafish, with dramatic downstream effects on pigment cells and pigmentation, as well as on the Notch signaling pathway (Akanuma et al. 2007; Nguyen et al. 2010). Perhaps unsurprisingly then, we found that *notch1*, a well-known member of the Notch Signaling Pathway, was differentially expressed between color morphs. Mutations in this gene are known to affect skin, hair and eye pigmentation in humans through effects on melanocyte stem cells (Schouwey and Beermann 2008). This indicates that *notch1* is a good candidate gene for pattern development in poison frogs.

A number of other melanogenesis-related genes were found to be differentially expressed between morphs, such as *brca1*. Mice with a homozygous mutation of the tumor suppressing *brca1* gene show altered coat coloration, often producing a piebald appearance (Ludwig et al. 2001). The precise mechanism behind this is ambiguous, and it may involve either *mitf* or *p53 (Beuret et al. 2011; Tonks et al. 2012). Bmpr1b* is a bone morphogenic protein which is known to inhibit melanogenesis; when *bmpr1b* is downregulated via UV exposure it enhances melanin production and leads to darker pigmentation (Yaar et al. 2006). Some of the other genes (e.g. *mlph*, or melanophilin) show the same pattern of expression across morphs as *lef1*, suggesting that multiple genes may contribute to the difference between lighter and darker background coloration in this species. The product of the melanophilin gene forms a complex that combines with two other proteins and binds melanosomes to the cell cytoskeleton, facilitating melanosome transport within the cell. Variants of this gene are associated with “diluted”, or lighter-colored, melanism in a number of vertebrates (Cirera et al. 2013). Similarly, the *mreg* (melanoregulin) gene product functions in melanosome transport and hence is intimately involved in pigmentation (Wu et al. 2012). Mutations at this locus cause “dilute” pigmentation phenotypes in mice.

In summary, we have found a number of differentially expressed genes that influence melanic coloration which seem to be important between color morphs with a true, black background pattern versus those with a more dilute, brown colored background pattern. Our results parallel similar findings in *Oophaga histrionica*, a species of poison frog in which mutations in the *mc1r* gene affecting melanogenesis have produced a lighter, more brownish background in some populations (Posso-Terranova and Andrés 2017). In addition to *mc1r*, we have identified a suite of genes with the same expression pattern that are ultimately influenced by *mc1r* activity; many of these genes have been linked to lighter phenotypes in other taxa.

### Purine synthesis and iridophore genes

The bright coloration of *D. auratus* is confined to the green-blue part of the visual spectrum (with the exception of some brownish-white varieties) in most populations, and thus iridophores are likely to play a role in the color variation displayed across different populations of this species. Higdon et al. (2013) identified a variety of genes that are components of the guanine synthesis pathway and show enriched expression in zebrafish iridophores. A number of these genes (*hprt1*, *ak5*, *dera*, *ednrb2*, *gas1*, *ikpkg*, *atic*, *airc*, *prtfdc1*) were differentially expressed between the different morphs of *D. auratus* investigated here (Figure 8). The *gart* gene codes for a tri-function enzyme that catalyzes three key steps in the *de novo* purine synthesis pathway (Ng et al. 2009). This locus has been associated with critical mutations affecting all three types of chromatophores in zebrafish, through effects on the synthesis of guanine (iridophores), sepiapterin (xanthophores) and melanin (melanocytes)(Ng et al. 2009). Zebrafish mutants at this locus can show dramatically reduced numbers of iridophores, resulting in a lighter, or less saturated color phenotype. Similarly, the *airc* gene plays a critical role in guanine synthesis, and yeast with mutations in this gene leading to aberrant forms of the transcribed protein are unable to synthesize adenine and accumulate a visible red pigment (Tolstorukov and Efremov 1984; Sychrova et al. 1999). Similarly, the *mthfd* (methylenetetrahydrofolate dehydrogenase, cyclohydrolase and formyltetrahydrofolate synthetase 1) gene also affects the *de novo* purine synthesis pathway (Christensen et al. 2013). The genes *airc*, *gart*, and *mthfd* had similar expression patterns and were very lowly expressed in the mostly green microspot population. The gene *prtfdc1* is highly expressed in iridophores, and encodes an enzyme which catalyzes the final step of guanine synthesis (Higdon et al. 2013); *prtfdc1* had very low expression in the dark blue-black morph, which may be an indication that it plays a role in the reflectance from iridophores. Further, *prtfdc1* was highly expressed in the San Felix and super blue morphs, both of which have visible small white ‘sparkles’ on the skin which are likely produced by the iridophores.

How the guanine platelets are formed in iridophores remains an open question. Higdon et al. (2013) proposed that ADP Ribosylation Factors (ARFs) and Rab GTPases are likely to play crucial roles in this context. ARFs are a family of ras-related GTPases that control transport through membranes and organelle structure. We identified one ARF protein (*arf6*) and two ARF activating proteins (*arfgap1* and *arfgap2*) that were differentially expressed across the *D. auratus* morphs. We also identified four different Rab GTPases as differentially expressed (*rab1a*, *rab3c*, *rab3d*, *rab7a*). Mutations at the *rabggta* (Rab geranylgeranyl transferase, a subunit) locus cause abnormal pigment phenotypes in mice (e.g. “gunmetal”), are known to affect the guanine synthesis pathway (Gene et al. 2001), and are similarly differentially expressed between color morphs in our dataset. These genes are likely candidates to affect coloration in *Dendrobates auratus* given that both the green and blue pattern elements are probably iridophore-dependent colors.

### Pteridine synthesis

A number of the genes identified as differentially expressed are involved in copper metabolism (*sdhaf2*, *atox1*, *atp7b*). Copper serves as a key cofactor for tyrosinase in the melanogenesis pathway and defects in copper transport profoundly affect pigmentation (Setty et al. 2008). Another gene, the xanthine hydrogenase (*xdh*) locus, was also found to be differentially expressed between morphs, and this gene, which is involved in the oxidative metabolism of purines, affects both the guanine and pteridine synthesis pathways. Additionally, it has been shown to be critically important in the production of color morphs in the axolotl. When *xdh* was experimentally inhibited axolotls had reduced quantities of a number of pterins, and also had a dramatic difference in color phenotype with *xdh*-inhibited individuals showing a ‘melanoid’ (black) appearance (Thorsteinsdottir and Frost 1986). Furthermore, *xdh* deficient frogs show a blue coloration in typically green species (Frost 1978; Frost and Bagnara 1979). We note here that one *xdh* transcript showed little (one individual) or no (2 individuals) expression in the bluest morph (blue-black). Similarly, when pigments contained in the xanthophores that absorb blue light are removed, this can lead to blue skin (Bagnara et al. 2007). We also found another gene involved in pteridine synthesis, *qdpr* (quinoid dihydropteridine reductase), was only expressed in the populations with a lighter blue or green coloration. Mutations in this gene result in altered patterns of pteridine (e.g. sepiapterin) accumulation (Ponzone et al. 2004). We believe that *xdh* and *qdpr* are good candidates for variability in coloration in poison frogs.

### Novel candidate genes for coloration

In addition to those genes that have previously been linked to coloration which we have identified in our study, we would like to propose several others as candidate color genes, based on their expression patterns in our data. Although most research on blue coloration focuses on light reflecting from iridophores, this has generally not been explicitly tested and there is some evidence that blue colors may arise through different mechanisms (reviewed in (Bagnara et al. 2007). In particular, there is evidence that blue in amphibians can come from the collagen matrix in the skin, as grafts in which chromatophores failed to thrive show a blue coloration (Bagnara et al. 2007). Furthermore, keratinocytes surround melanocytes, and they play a key role in melanosome transfer (Ando et al. 2012). In light of this evidence, we propose a number of keratinocyte and collagen genes which are differentially expressed in our dataset as further candidate genes for coloration. Amongst these are *krt12*, and *krt18*, *col1a1*, *col5a1*, and *col14a1*. These genes, and those like them, may be playing a critical role in coloration in these frogs.

### Conclusion

The mechanisms that produce variation in coloration in both amphibians and aposematic species are poorly characterized, particularly in an evolutionary context. Here we have taken a transcriptomics-based approach to elucidating the genetic mechanisms underlying color and pattern development in a poison frog. We found evidence that genes characterizing the melanin and iridophore pathways are likely the primary contributors to color and pattern differences in this aposematic species. Additionally, a handful of genes which contribute to the pteridine pathway seem to be playing a role in differential color production as well. However, the specific mechanisms by which these genes work, as well as how they interact to produce color phenotypes, remains an outstanding issue given the complex nature of each of these pathways. Still, our data indicate that genes involved at every step along the melanin and iridophore pathways from chromatophore production, through pigmentation production and deposition, influence differences in coloration between these morphs. These results make sense in the context of the overall color and pattern of these frogs, and provide a number of promising starting points for future investigations of the molecular, cellular and physiological mechanisms underlying coloration in amphibians.

## Acknowledgments

Animal care and use for this research was approved by East Carolina University’s IACUC (AUP #D281). Funding for this project was provided by NSF DEB 165536 and an East Carolina University Thomas Harriot College of Arts and Sciences Advancement Council Distinguished Professorship to K Summers. We thank Evan Twomey for insightful comments on this manuscript.

